# When does spatial diversification usefully maximise the durability of crop disease resistance?

**DOI:** 10.1101/540013

**Authors:** Benjamin Watkinson-Powell, Christopher A. Gilligan, Nik J. Cunniffe

## Abstract

Maximising the durability of crop disease resistance genes in the face of pathogen evolution is a major challenge in modern agricultural epidemiology. Spatial diversification in the deployment of resistance genes, where susceptible and resistant fields are more closely intermixed, is predicted to drive lower epidemic intensities over evolutionary timescales. This is due to an increase in the strength of dilution effects, caused by pathogen inoculum challenging host tissue to which it is not well-specialised. The factors that interact with and determine the magnitude of this spatial effect are not currently well understood however, leading to uncertainty over the pathosystems where such a strategy is most likely to be cost-effective. We model the effect on landscape scale disease dynamics of spatial heterogeneity in the arrangement of fields planted with either susceptible or resistant cultivars, and the way in which this effect depends on the parameters governing the pathosystem of interest. Our multi-season semi-discrete epidemiological model tracks spatial spread of wild-type and resistance breaking pathogen strains, and incorporates a localised reservoir of inoculum, as well as the effects of within and between field transmission. The pathogen dispersal characteristics, any fitness cost(s) of the resistance breaking trait, the efficacy of host resistance, and the length of the timeframe of interest, all influence the strength of the spatial diversification effect. These interactions, which are often complex and non-linear in nature, produce substantial variation in the predicted yield gain from the use of a spatial diversification strategy. This in turn allows us to make general predictions of the types of system for which spatial diversification is most likely to be cost-effective, paving the way for potential economic modelling and pathosystem specific evaluation. These results highlight the importance of studying the effect of genetics on landscape scale spatial dynamics within host-pathogen disease systems.

## 2 Introduction

The evolution of pathogen populations in response to the use of resistant crop cultivars is a major area of concern for global food production. There is also environmental concern, as a lower efficacy of resistant crops generally leads to a greater reliance on chemical pesticides, with potential negative impacts on non-agricultural ecosystems. Increasing the durability of crop disease resistance, defined as the period of time that resistance remains effective with its widespread cultivation, is therefore an important focus for plant breeders and epidemiologists (Johnson, 1979). Resistances can break down in a matter of years or even months, with resistances to fungal and bacterial diseases generally breaking down faster than those for viruses, in part due to the often lower mutational fitness costs in cellular pathogens (Garcia-Arenal and McDonald, 2003). The resultant effect of this breakdown can be a significant reduction in crop yields, underlining the need to understand how host resistances can be effectively deployed.

The interaction between plant disease resistance and corresponding pathogen virulence, defined here as the qualitative ability of a pathogen to infect a host, often takes the form of a gene-for-gene relationship (Flor, 1971). This genetic system is defined by a single *R* gene in the host that recognises, and provides qualitative resistance to, a single corresponding avirulence gene in the pathogen. Often this avirulence gene has an important role as an effector in the mechanism of infection (Cui et al., 2015). Mutations in the avirulence gene can cause it to evade the corresponding *R* gene and become ‘resistance breaking’ or ‘virulent’ (Flor, 1971). However, there is substantial evidence that resistance breaking traits in plant pathogens have high reproductive fitness costs, with examples in plant viruses (Fraile et al., 2011), and basidomycetes (Thrall and Burdon, 2003, Bruns et al., 2014). These costs have the potential to prevent the resistance breaking strain from completely dominating when a mixture of host cultivars are deployed. On a fully susceptible host, which can be infected by both wild-type and resistance breaking pathogen strains, the costs of the resistance breaking trait cause it to experience negative selective pressure. In the absence of immigration or new mutation, this would ultimately lead to loss of the resistance breaking strain from the pathogen population. Such loss of unnecessary virulence is seen in field experimental evolution studies, where the lack of positive selection can result in a reduced frequency of individual resistance breaking pathogen strains (Bousset et al., 2018). On the other hand, mutation to virulence can occur quite frequently, with wild-type viruses for example becoming resistance breaking with as few as one or two nucleotide substitutions in the genes affecting avirulence (Harrison, 2002). This can lead to resistance breaking strains persisting at very low background frequencies in pathogen populations, even in the absence of selection from the corresponding resistant hosts (Stam and McDonald, 2018).

As with any process which occurs over relatively long periods of time and over large spatial scales, mathematical modelling is a useful tool for examining the factors affecting resistance durability. In recent years, modelling in this area has integrated population genetics and epidemiology to consider durability largely in terms of the overall amount of host plant material infected in a given time period. van den Bosch and Gilligan (2003) introduced the number of additional uninfected host growth days as a measure of resistance durability, moving the focus away from simply looking at pathogen genotype frequencies as had often been done previously (Brown, 1995, Lannou and Mundt, 1996). The work by van den Bosch and Gilligan (2003) was further developed by Fabre et al. (2012), who represented the seasonally disturbed nature of the agricultural environment with a semi-discrete model (Mailleret and Lemesle, 2009). In the Fabre et al. (2012) model, disease spread occurs in continuous time during cropping seasons but is reset upon harvesting, with dynamics in subsequent years being affected by a reservoir component which allows the pathogen to overwinter between discrete seasons (Burdon and Thrall, 2008). Fabre et al. (2012) also implicitly characterised landscape spatial structure by modifying the relative contributions of within field, between field, and reservoir driven infection. This characterisation captures the overall degree of connectedness between different fields, along with the relative importance of ongoing primary infection from the reservoir component, but takes no account of explicit spatial effects such as dispersal distances or the scale or pattern of spatial heterogeneity. Results from the study by Fabre et al. (2012) highlighted the impact of resistance gene choice on durability, in terms of how easily it can be overcome by mutations in the pathogen avirulence gene, and with what associated fitness costs.

It is known from both theoretical and experimental studies that the mixing of cultivars with differing resistance properties can reduce the rate of disease spread in single and multi-pathogen strain systems (Mundt, 2002, Zhan and McDonald, 2013). This can occur through dilution effects, where some of the force of infection from a given pathogen strain is wasted due to its reduced ability, or even inability, to infect the portions of the host population on which it is not specialised (Mundt, 2002). It has been found in some theoretical model pathosystems that within-field cultivar mixing is more effective at controlling disease spread than planting some fields wholly with susceptible plants and others with resistant plants (Skelsey et al., 2010). There are however frequent complications associated with the use of within-field mixtures, including phenotypic differences between host varieties, such as in harvest dates, along with the perceived need for product purity (Burdon et al., 2016). These factors, which in part explain the limited use of mixtures in agriculture in the developed world, highlight the need to understand the effects on disease dynamics of employing host diversification at larger scales than within a single field, i.e. at the landscape or between field scale.

The number of studies looking at the effects of landscape scale host diversification combined with long range pathogen dispersal is limited, no doubt held back by the largely field scale nature of empirical work in this area (Plantegenest et al., 2007). This topic has received a degree of attention in theoretical studies however, with Skelsey et al. (2010) finding that the clustering of potato cultivation reduced the spread of late blight between clusters, but increased overall epidemic intensity due to higher spread within clusters. It has been posited that this scenario can potentially create a trade-off between within and between patch dispersal, resulting in maximised dispersal at intermediate scales of spatial clustering (Skelsey et al., 2013). Spatial structure is also relevant over longer timescales, with Papaïx et al. (2013) showing, in a general theoretical metapopulation study, that spatial clustering of habitat patches facilitates specialisation within a population, in addition to driving increased evolutionary speeds.

One component of landscape scale spatial structure that can potentially be optimised for disease control and resistance durability is the scale of spatial heterogeneity in the deployment of different host cultivars. The mechanism behind the efficacy of mixtures suggests that mixing host genotypes at smaller scales of spatial heterogeneity, and thereby creating smaller genotype unit areas (GUAs), would benefit disease control by decreasing connectivity between patches of the same host cultivar (Mundt, 2002). A modelling study by Papaïx et al. (2014), concerning a single pathogen strain, found similar benefits to mixing crop genotypes at smaller spatial scales when using major genes conferring complete resistance. However these authors also found that for some levels of incomplete resistance, small patches of partially resistant crop could act as sinks for nearby pathogen populations in susceptible patches, and thereby act as stepping stones to increase overall disease incidence in the landscape with greater field mixing.

Some studies have looked more explicitly at resistance durability against multiple pathogen strains over evolutionary timescales. Sapoukhina et al. (2009) showed, using a reaction-diffusion model, that random mixtures in the host landscape provide greater long term disease suppression compared with patchy mixtures at larger scales of spatial heterogeneity. This is supported by a recent modelling study by Papaïx et al. (2018), who showed that low levels of spatial aggregation, in a mixed landscape of fields planted with either susceptible or resistant cultivars, reduced epidemic intensities over both short term epidemiological timescales and at the long term evolutionary equilibrium. A field experimental evolution study by Bousset et al. (2018) provides a degree of empirical support for this general theory, with results suggesting that higher ‘genetic connectivity’ within a host population facilitates higher levels of infection. Arguably, a potential weakness of this study was the implicit representation of ‘genetic connectivity’ by greater experimental inoculation of host variety patches by their specialist pathogen strains, which means that it did not truly demonstrate a landscape scale spatial effect.

A number of recent theoretical studies have begun comparing and contrasting the various available strategies for the optimal deployment of resistance genes, such as using mosaics (between field spatial diversification), mixtures, rotations and pyramids (Fabre et al., 2015, Djidjou-Demasse et al., 2017, Rimbaud et al., 2018). We believe however that the specific role of spatial diversification and the dynamics which affect this strategy are not currently well understood. While existing studies consistently point to smaller scales of spatial heterogeneity being optimal for durable and effective disease control in multi-strain systems, there is generally little investigation of the factors that influence the strength of this spatial effect. In order for such spatial strategies to be employed in commercial agriculture, they will need to be cost effective, in that the benefit to resistance durability of planting fields of different cultivars at smaller scales of spatial heterogeneity must offset the likely increased financial cost and operational difficulty of farming in this manner. The principal aim of this study is therefore to examine the factors interacting with, and influencing the strength of, any such spatial effect. These factors include the dispersal characteristics of the pathogen, the fitness costs associated with the resistance breaking trait, the efficacy of the host resistance gene, and the length of the timescale of interest.

## 3 Materials and Methods

We extend the model used by Fabre et al. (2012) to explicitly include spatial structure and different pathogen strains. Our *SI* (Susceptible, Infected) model tracks two pathogen strains, a ‘wild-type’ (*wt*) and a ‘resistance breaking’ (*rb*) genotype (the principal variables and parameters used in this model are summarised in table 1). The underlying host landscape consists of a number (*n*_*f*_ = 100) of cultivated fields, each with a constant number of plants/plant tissue units (*n*_*p*_ = 1000) of either a susceptible (*S*) or resistant (*R*) cultivar type (note that the use of continuous state variables combined with the method of epidemic parametrisation makes the results ultimately independent of *n*_*p*_). The proportion of resistant fields (the cropping ratio) is set evenly with that of susceptible fields at *ϕ* = 0.5.

**Table 1:** Parameters and variables used in the mathematical model

| Symbol | Parameter/Variable Description | Constraints/Values |
| --- | --- | --- |
| $I_{wt,x,y}$ | Number of plants in field $x$ and season $y$ infected by the $wt$ strain | |
| $I_{rb,x,y}$ | Number of plants in field $x$ and season $y$ infected by the $rb$ strain | |
| $n_f$ | Number of fields | 100 |
| $n_p$ | Number of plants/plant tissue units per field | 1000 |
| $\phi$ | Cropping ratio (proportion of resistant fields) | 0.5 |
| $n_d$ | Number of days in a season | 120 |
| $n_y$ | Number of seasons | $1 \leq n_y \leq 80$ |
| $\theta$ | Background equilibrium frequency of $rb$ strain on $S$ host | 0.01 |
| $\delta$ | Fitness cost of the $rb$ trait | $0 \leq \delta \leq 1$ |
| $\gamma$ | Susceptibility of the $R$ host to the $wt$ strain | $0 \leq \gamma \leq 1$ |
| $K[z, x]$ | 2D normalised dispersal kernel coupling field $z$ to field $x$ | $z \neq x, K = \frac{\eta^2}{2\pi} e^{-\eta d}$ |
| $\eta$ | Dispersal kernel parameter | 3 levels: 1, 2, 3 |
| $A_0$ | Baseline AUDPC for one season in a fully susceptible landscape | $A_0 = \sum_{x=1}^{n_f} (\int_0^{n_d} I_{S,x,y}(t) dt)$ |
| $\beta_F$ | Within field infection rate | (see text and SI) |
| $\beta_C$ | Between field infection rate | (see text and SI) |
| $\alpha_E$ | Baseline rate of infection from the reservoir | (see text and SI) |
| $\alpha_{wt,y}$ | Rate of $wt$ infection from the reservoir | |
| $\alpha_{rb,y}$ | Rate of $rb$ infection from the reservoir | |
| $A_{wt,x,y-1}$ | AUDPC for the $wt$ epidemic in field $x$ in the previous season | $\int_0^{n_d} I_{wt,x,y-1}(t) dt$ |
| $A_{rb,x,y-1}$ | AUDPC for the $rb$ epidemic in field $x$ in the previous season | $\int_0^{n_d} I_{rb,x,y-1}(t) dt$ |

Resistance acts in a gene-for-gene system where the *wt* strain can freely infect *S* fields, but has reduced fitness (that may be zero) when infecting *R* fields. The *rb* strain has equal infective ability on both host genotypes, but may have a reproductive fitness cost (*δ*) (expressed on both host genotypes) associated with its resistance breaking trait. The focus of our model on disease spread at the landscape scale, driven by long range pathogen dispersal, makes it appropriate for application to any gene-for-gene crop disease system with a foliar wind dispersed pathogen (such as many rusts or powdery mildews).

The overall number of infected plants in each field is simulated for *n*_*d*_ = 120 days over *n*_*y*_ seasons in a semi-discrete modelling approach. In this, continuous time dynamics in ordinary differential equations (ODEs) are used for the within season component, and discrete dynamics are used for the pathogen infecting and overwintering in the reservoirs (Mailleret and Lemesle, 2009). The reservoir components are localised to each field, and represent plants found in field margins and hedgerows, as well as crop volunteers, that act as alternative hosts for the pathogen. In all simulations, *I*_*wt,x,y*_(0) = *I*_*rb,x,y*(0)_ = 0 (i.e. the number of infected plants within fields is set to zero at the beginning of each season), with all epidemics started by primary infection from the reservoir of inoculum.

The total area under the disease progress curve (AUDPC) over the *n*_*y*_ seasons is used as the measure of epidemic intensity (Madden et al., 2007). As a measure of resistance durability this is equivalent to the number of uninfected host growth days used by van den Bosch and Gilligan (2003). The AUDPC is normalised to a value between 0 and 1 to obtain the average proportion of plants infected across the sequence of epidemic seasons (epidemic intensity = AUDPC/(*n*_*f*_ *n*_*p*_*n*_*y*_*n*_*d*_)). Healthy plants can become infected through three alternative routes: from infected plants in the same field at rate *β*_*F*_, from infected plants in other fields at rate *β*_*C*_, and from the reservoir of inoculum at rates *α*_*wt,y*_ or *α*_*rb,y*_ in season *y* for the *wt* and *rb* pathogen strains respectively. The pathogen population size in the reservoir is not explicitly modelled but is represented by these rate parameters (*α*_*wt,y*_ and *α*_*rb,y*_) by scaling the baseline rate of infection from the reservoir component *α*_*E*_.

The values of the transmission parameters *β*_*F*_, *β*_*C*_ and *α*_*E*_ were optimised by following Fabre et al. (2012) to calculate the relative contributions of the three infection routes to, and the overall intensity of, a baseline epidemic scenario (see Supporting Information Notes S1). The version of the model used for this optimisation uses a landscape with susceptible fields only (*ϕ* = 0). The values of the transmission parameters are such that in this case the three transmission routes have an equal contribution to maintaining the epidemic, and the mean proportion of plants infected during a season is 0.5.

The fields are located within a square landscape of 10×10 arbitrary distance units, where the distances (*d*) between each pairwise combination of fields are calculated and used in a normalised negative exponential dispersal kernel (of the form 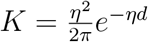 with mean dispersal distance 1/*η*) to calculate each pairwise strength of spatial coupling. Each simulation was repeated 100 times, with different random field locations in each replicate.

To explore the effect of the scale of spatial heterogeneity in the arrangement of *S* and *R* fields on epidemic intensities, the landscape area is arranged (with cropping ratio *ϕ* = 0.5) into a number of patches. Each patch only contains fields of a single host genotype (*S* or *R*). At the largest scale the landscape is split in half, with all the *S* fields in one half and all the *R* fields in the other. At the smallest scale the landscape is split into 64 patches with alternating host genotypes. Four intermediate scales are also used, for a total of six landscape templates with different patch sizes and scales of spatial heterogeneity (table 2). We characterise the scale of spatial heterogeneity via the interior edge/area (E/A) ratio of each landscape (see Supporting Information Notes S2). This metric captures the degree of contact between the two different host patch types, relative to the overall size of the landscape. It serves as an effective proxy for the overall proximity of the two host field types to each other, and can also be applied to landscapes with irregular patch shapes, variable patch sizes, and different total areas. The pattern of host genotype patches is used as a template to place the two types of field, using random coordinates, in alternating patches within the landscape (Fig. 1).

**Figure 1:**
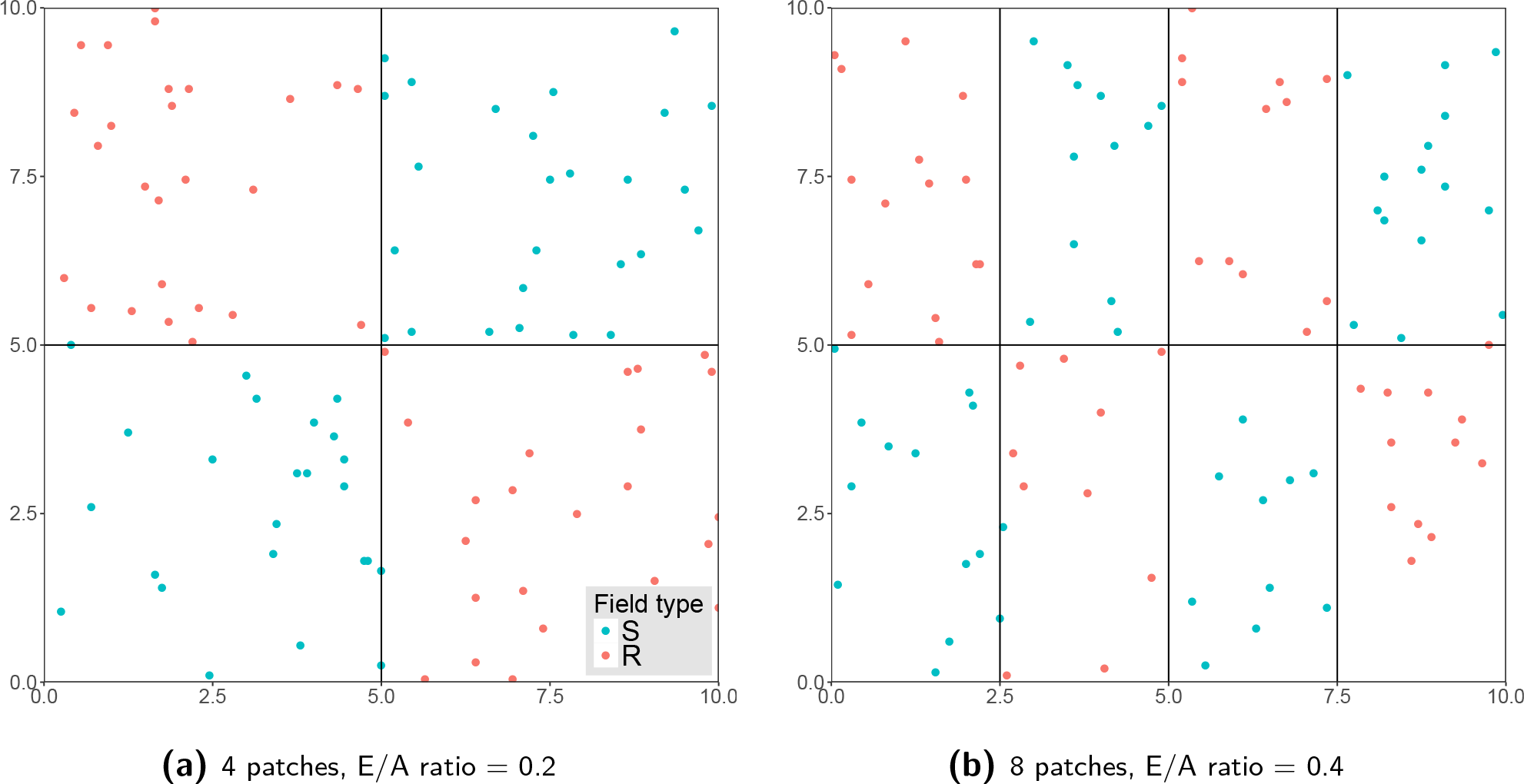
Example random field arrangements for two patch templates within the agricultural landscape.

**Table 2:**
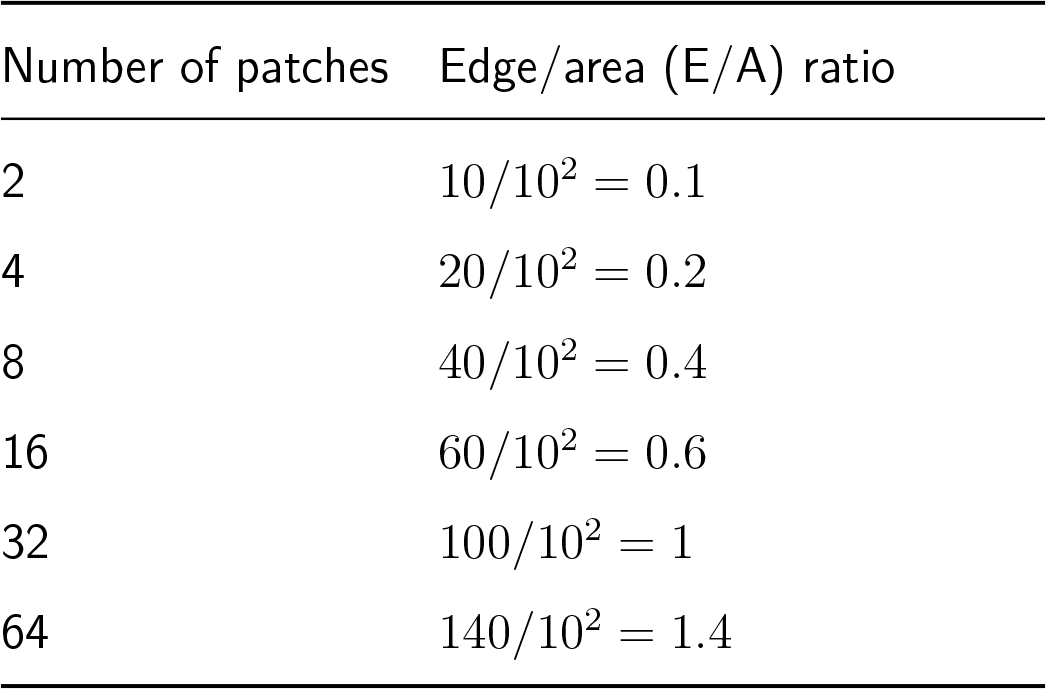
The edge/area ratios of the landscapes at each of the six scales of spatial heterogeneity (see also Fig. 1 and Supporting Information Notes S2).

The spread of the two pathogen strains in a given field within a season follows the general pseudo-equation form:

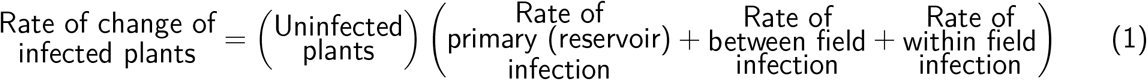

and is described by the deterministic ODE system:

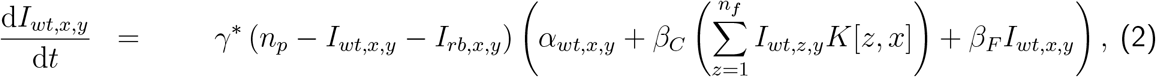

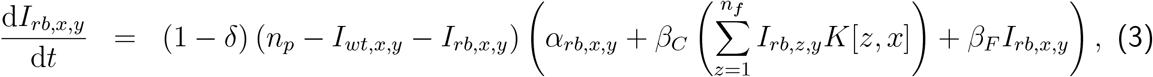

in which

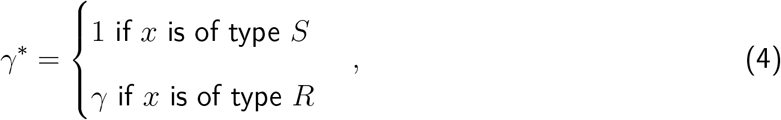

in Eqn (2).

In Eqns (2) and (3), *x* indicates variables pertaining to a particular field, *K*[*z, x*] is the dispersal kernel coupling field *z* to field *x*, and *z* ≠ *x*. Here *I*_*wt,x,y*_ is the number of plants infected by the *wt* strain while *I*_*rb,x,y*_ is the number infected by the *rb* strain (both being in field *x* and in season *y*). The cost of the resistance breaking trait is *δ*, which is assumed act on the rate of sporulation on both the primary and reservoir host, while *γ* is the susceptibility of the *R* host to the *wt* strain. If resistance to the *wt* strain is complete *γ* = 0, meaning that the *wt* epidemic in an *R* field (Eqn (2) where *γ*\* = *γ*) does not occur.

The terms *α*_*wt,x,y*_ and *α*_*rb,x,y*_ represent the specific infection rates of each pathogen genotype from the local reservoir, where the reservoir host is assumed to be selectively neutral to both pathogen genotypes. These rates are subject to change between seasons from initial values of *α*_*wt,x*,0_ = *α*_*E*_ and *α*_*rb,x*,0_ = *θα*_*E*_, where *θ* is the background equilibrium frequency of the *rb* genotype resulting from the mutation-selection balance in the absence of positive selection from the deployment of the resistant host (Fabre et al., 2012). The contributions of the reservoir components to the epidemic in season *y* are given by:

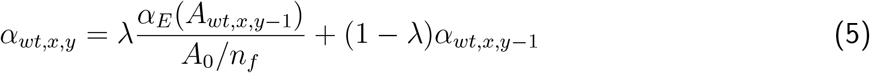

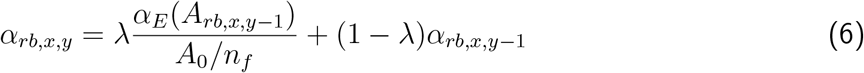

Here, 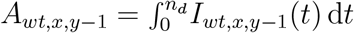 and 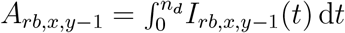, the AUDPCs for the epidemics in the previous season caused by the *wt* and *rb* pathogen genotypes respectively in a given individual field *x*. The baseline landscape AUDPC 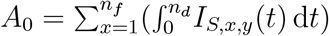, calculated from one season in the fully susceptible baseline model, is used to scale the previous season’s epidemic to measure the proportional reduction in epidemic intensity due to the presence of *R* fields.

The parameter *λ* ∈ [0, 1] characterises the reservoir. High values of *λ* indicate a rapidly changing reservoir with primary infection largely depending on the intensity of the previous season’s field epidemics. This scenario could potentially be caused by a short reservoir host lifespan, a high rate of spread within the reservoir, or a small reservoir size. Low values of *λ* indicate a more ‘stable’ reservoir, with a larger effect of older epidemics damping changes to *α*_*wt,x,y*_ and *α*_*rb,x,y*_. A value of *λ* = 0.5, equally weighting both terms in Eqns (5) and (6), is used here.

## 4 Results

An illustrative time series of seasonal disease progress curves for an example landscape with a high edge/area ratio is shown in Fig. 2. The average seasonal epidemic intensity (average proportion of plants infected throughout the epidemic) over a 40 season time period invariably decreases as the edge/area (E/A) ratio of the landscape is increased (Fig. 3). This is due to the larger dilution caused by greater mixing of the two host genotype field types at smaller scales of spatial heterogeneity. Epidemic intensity is initially measured here over a 40 season time period, to balance short term and long term evolutionary dynamics, although the effect of varying the time frame of interest is described in section 4.4. The reduction in epidemic intensities due to the planting of *S* and *R* fields at smaller scales of spatial heterogeneity within the landscape (higher E/A ratios) is variously referred to as the ‘spatial effect’, ‘spatial suppressive effect’ or ‘suppressive effect’ throughout the remainder of this article.

**Figure 2:**
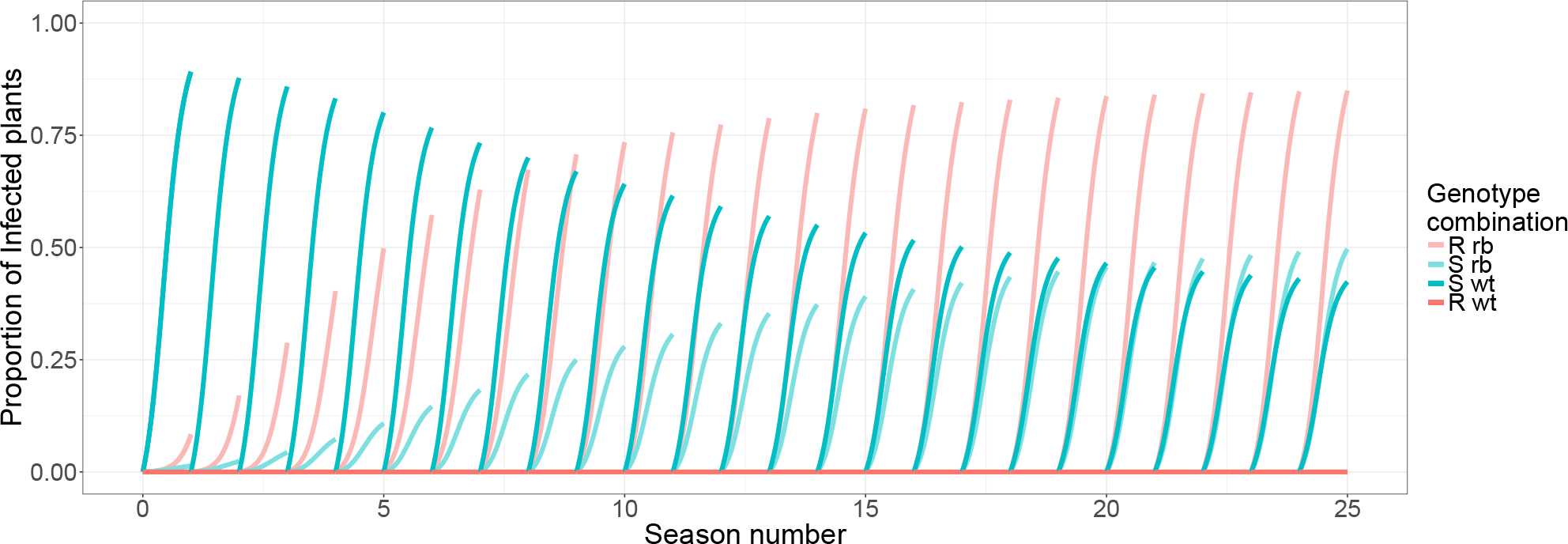
Seasonal disease progress curves for each infection class (genotype combination) in an example landscape simulation replicate. The intensity of disease changes between seasons due to the changing contribution of primary infection from the local reservoirs, which in turn is based on the local epidemic intensities in the previous season. The two field types in the landscape are highly mixed (edge/area ratio = 1.4), the cost of the *rb* trait *δ* = 0.3 and the *R* host is completely resistant to the *wt* strain *γ* = 0. Only 25 seasons are shown here so that the individual disease progress curves can see clearly seen.

**Figure 3:**
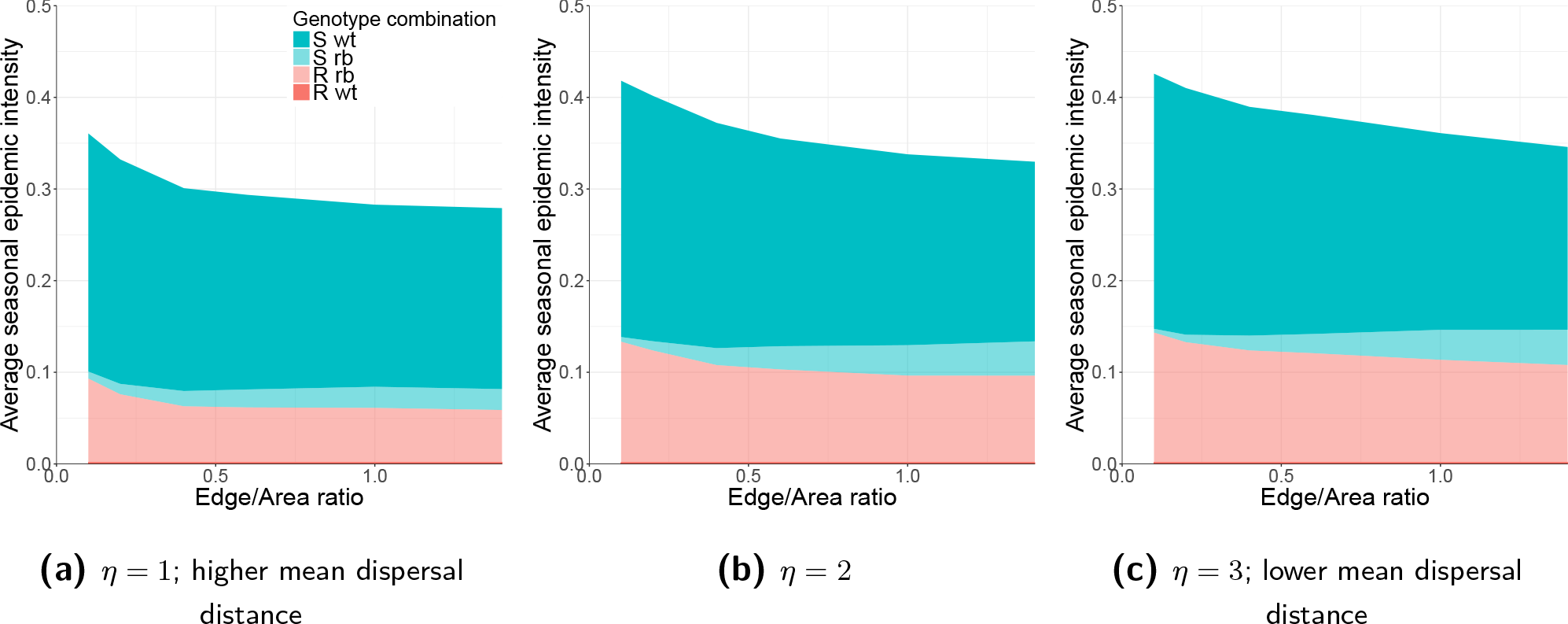
The effect of the kernel parameter *η* on the relationship between landscape edge/area ratio and epidemic intensities. **Epidemic intensities over evolutionary timescales are reduced at smaller scales of spatial heterogeneity. The spatial scale at which a spatial strategy is effective depends on the scale of pathogen dispersal.** For all simulations presented here, the number of seasons *n*_*y*_ = 40, the cost of the *rb* trait *δ* = 0.3 and the *R* host is completely resistant to the *wt* strain *γ* = 0.

The *wt* epidemic is suppressed at higher E/A ratios (Fig. 3) due to the smaller spatial grain causing an increase in the proportion of dispersed *wt* inoculum from *S* fields that is wasted as it lands on, but is unable to infect, *R* fields. The *rb* epidemic in *R* fields meanwhile is suppressed by a corresponding process in which there is an increase in the proportion of *rb* inoculum that lands on *S* fields. The *rb* inoculum is in direct competition with *wt* inoculum for uninfected host tissue within *S* fields, resulting in a lower intensity *rb* epidemic than would take place in an *R* field. The consequent *rb* genotype dispersal from these *S* fields back onto the nearby *R* fields therefore has a lower force of infection than in *R* field to *R* field transmission over the same distance. The reduction in the intensity of the *rb* epidemic on *R* fields is compensated to a certain extent by the increase in the frequency of the *rb* genotype on *S* fields as the two field genotypes are more closely mixed together in space. Whether this compensation ultimately increases or decreases the intensity of the overall landscape *rb* epidemic, as the E/A ratio is increased, depends upon the genetic parameters (Fig. 4).

**Figure 4:**
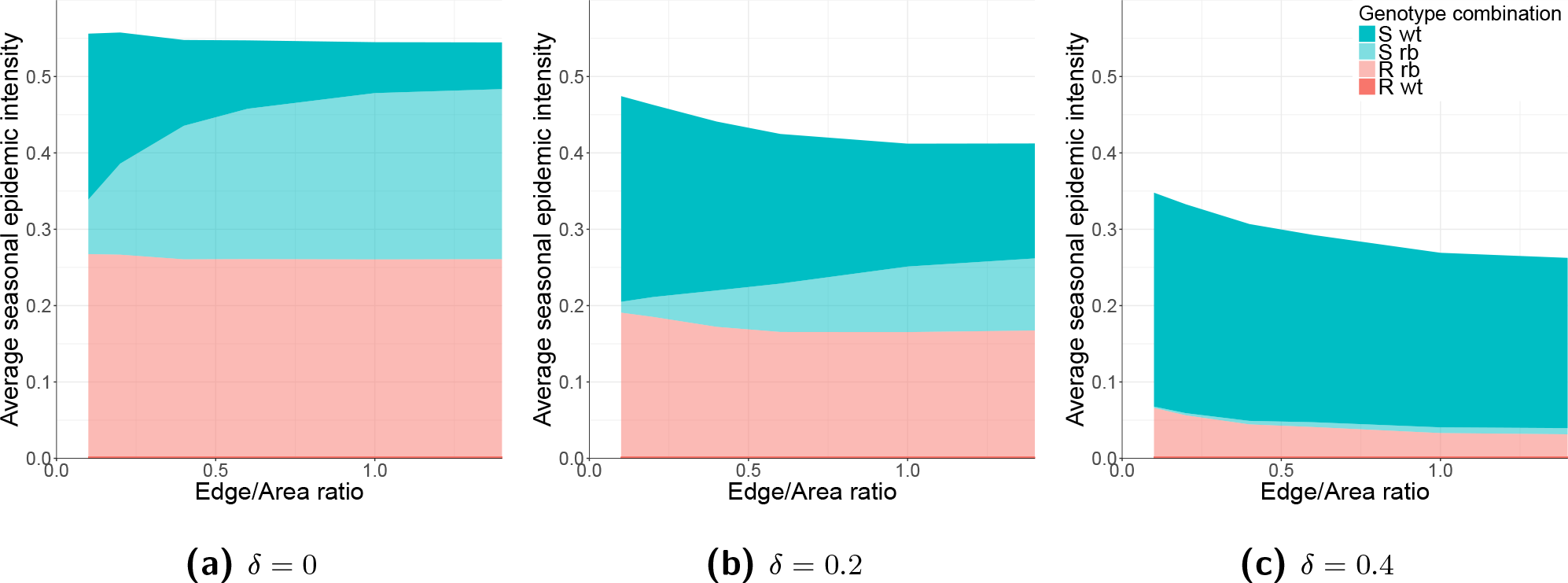
The effect of the fitness cost of the resistance breaking trait (*δ*) on the relationship between landscape edge/area ratio and epidemic intensities. **The strength of the spatial effect increases as the fitness cost of the resistance breaking trait is increased from low values. The pathogen genotype frequencies depend on the scale of spatial heterogeneity.** For all simulations presented here, the number of seasons *n*_*y*_ = 40, the *R* host is completely resistant to the *wt* strain *γ* = 0 and the kernel parameter *η* = 2.

### 4.1 Effect of the kernel parameter *η*

The gradient of the spatial suppressive effect (the rate of change in the response of average seasonal epidemic intensity to E/A ratio) decreases as the E/A ratio is increased (Fig. 3). The rate of gradient change at low E/A ratios is faster however with a higher mean dispersal distance (flatter dispersal kernel). This washes out and limits the strength of the spatial effect at smaller scales of spatial heterogeneity, where the high mean dispersal distance of the pathogen limits the impact of any further decrease in the scale of spatial heterogeneity. In general, this means that in the reverse direction, as the mean dispersal distance decreases, the spatial suppressive effect is relevant over a larger range of E/A ratios.

### 4.2 Effect of the cost of the rb trait *δ*

The overall size of the spatial effect increases as the cost of the *rb* trait *δ* is increased from 0 to 0.4 (Fig. 4). Furthermore, the extent to which the pathogen genotype frequencies on the *S* host change, as we move from larger to smaller scales of spatial heterogeneity, also depends on the cost of the *rb* trait. When *δ* = 0 (Fig. 4a) the *rb* genotype is able to take advantage of the greater proportion of host fields that it can infect, and the close proximity of the two field types at smaller spatial scales of heterogeneity, allowing it to outcompete the *wt* genotype. This replacement of pathogen genotypes at different E/A ratios is seen to a lesser extent when *δ* = 0.2 (Fig. 4b), and is almost absent when *δ* = 0.4 (Fig. 4c). As the fitness cost *δ* increases the *rb* genotype is unable to compete as effectively with the *wt* genotype on the *S* host at small scales of spatial heterogeneity. This is despite the close proximity of large numbers of *R* fields, which act as a major source of *rb* inoculum, to the *S* fields.

Plotting overall epidemic intensity against the full range of values for the fitness cost of the *rb* trait (*δ*), at both the low and high ends of the E/A ratio scale (i.e. for E/A= 0.1 and 1.4), allows these contrasting spatial scenarios to be compared (Fig. 5). For both E/A ratios, epidemic intensities decrease as *δ* is increased, up to *δ* = 0.6 where the *rb* trait is too expensive for that pathogen genotype to invade and there is no further effect of increasing *δ* (Fig. 5a). The variability and strength of the spatial suppressive effect can be ascertained by plotting the difference between the epidemic intensities for the two E/A ratio values (Fig. 5b). The strength of the suppressive effect increases from *δ* = 0 to 0.3, but decreases from *δ* = 0.4 to 0.6. A small suppressive effect is still seen at *δ* = 0 due to the small scale of spatial heterogeneity disrupting the transient *wt* epidemic, before the *wt* strain is outcompeted by the *rb* and reaches its near zero evolutionary equilibrium frequency.

**Figure 5:**
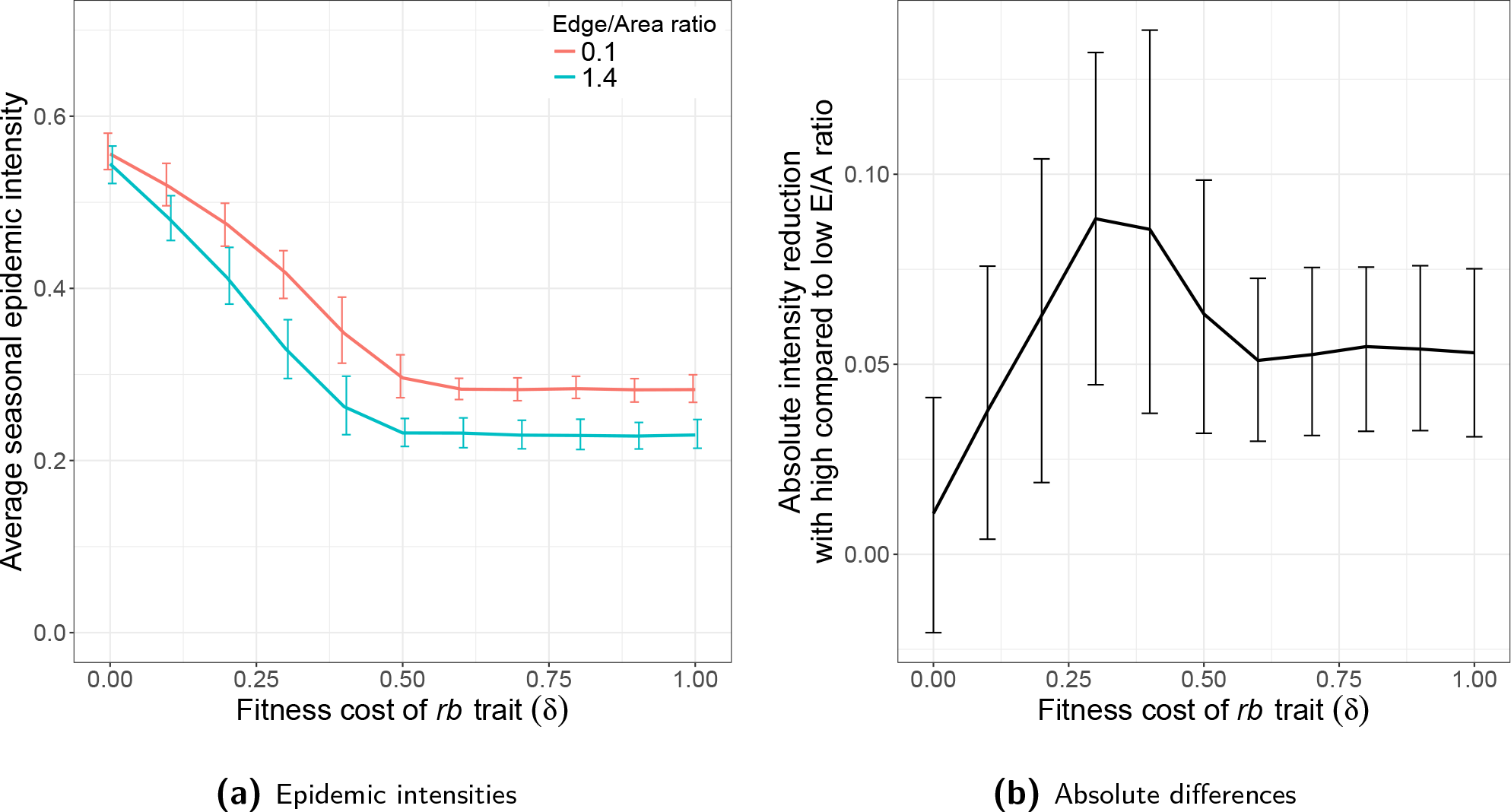
The effect of the fitness cost of the resistance breaking trait (*δ*) on the reduction in epidemic intensities from using a high compared to a low landscape edge/area ratio. **The strength of the spatial effect depends on the fitness cost of the resistance breaking trait, with a peak effect strength at an intermediate cost.** The average epidemic intensities resulting from the low and high ends of the E/A ratio scale are shown in (a), while the absolute differences between the results for these two E/A ratio values are shown in (b). Note that using the proportional differences in epidemic intensity produces a qualitatively similar pattern. Error bars show the 5th and 95th percentiles of the simulation replicates with stochastic landscape generation. For all simulations presented here, the number of seasons *n*_*y*_ = 40, the *R* host is completely resistant to the *wt* strain *γ* = 0 and the kernel parameter *η* = 2.

The initial increase in the strength of the suppressive effect is due to a steeper gradient of change, in the response of overall epidemic intensity to changes in *δ*, at small scales of spatial heterogeneity (E/A ratio = 1.4) (Fig. 5a). This steeper change with *δ* is in turn caused by rapidly changing *rb* dynamics on the *S* host, which are a larger driver of system sensitivity with greater field mixing (Supporting Information Fig. S2). Here, any increase in *δ* reduces the competitive ability of the *rb* genotype against the *wt* on susceptible hosts, which consequentially increases the amount of *rb* inoculum that is ‘wasted’ as it disperses onto these *S* hosts, thereby increasing the spatial effect strength.

The subsequent fall in the strength of the suppressive effect is correspondingly due to a steeper gradient of change with *δ* at large scales of spatial heterogeneity (E/A ratio = 0.1) (Fig. 5a). In this range of *δ* values there is an increased sensitivity to changes in *δ* of the *rb* epidemic on the *R* host, relative to that on the *S* host (Supporting Information Fig. S2). This is primarily because the *rb* genotype is already unable to compete effectively with the *wt* on the susceptible host in this range, and therefore does not respond to further changes in the fitness cost. These rapidly changing *R rb* dynamics are a larger driver of system sensitivity with less field mixing.

The value of *δ* for which the gradient of overall epidemic intensity change with *δ* is equal at both small and large scales of spatial heterogeneity (*δ* = 0.3 to 0.4), is the point of maximum spatial suppressive effect on epidemic intensities (i.e. the maximum distance between the curves). At this point the combined sensitivity effects, of *rb* dynamics on both *S* and *R* hosts to changes in *δ*, have the same net result at high and low landscape E/A ratios.

### 4.3 Partial resistance

Here, we relax the assumption that the *wt* strain cannot infect resistant hosts (Fig. 6). As *γ* is increased above 0, the *wt* genotype becomes able to infect the *R* host, and at higher frequencies with smaller scales of spatial heterogeneity (high E/A ratios) (Fig. 6a,b). The reduced efficacy of the resistance gene allows the *wt* strain to compete more effectively with the *rb* strain on the *R* host, particularly at high E/A ratios where the field types are more greatly mixed in space. The strength of the spatial suppressive effect, from using a high rather than a low E/A ratio, is shown in Fig. 6d. For an intermediate cost of the *rb* trait (*δ* = 0.3), the strength of the spatial suppressive effect decreases to zero as *γ* is increased from 0 to 0.6. This effect is due to a reduction in the proportion of *wt* inoculum that is ‘wasted’ in its increased dispersal onto *R* fields at smaller scales of spatial heterogeneity. When *γ* = 1 there is no effect of the scale of spatial heterogeneity, as the landscape is then made up of entirely susceptible hosts, and therefore the *wt* genotype is able to fully outcompete the *rb* genotype.

**Figure 6:**
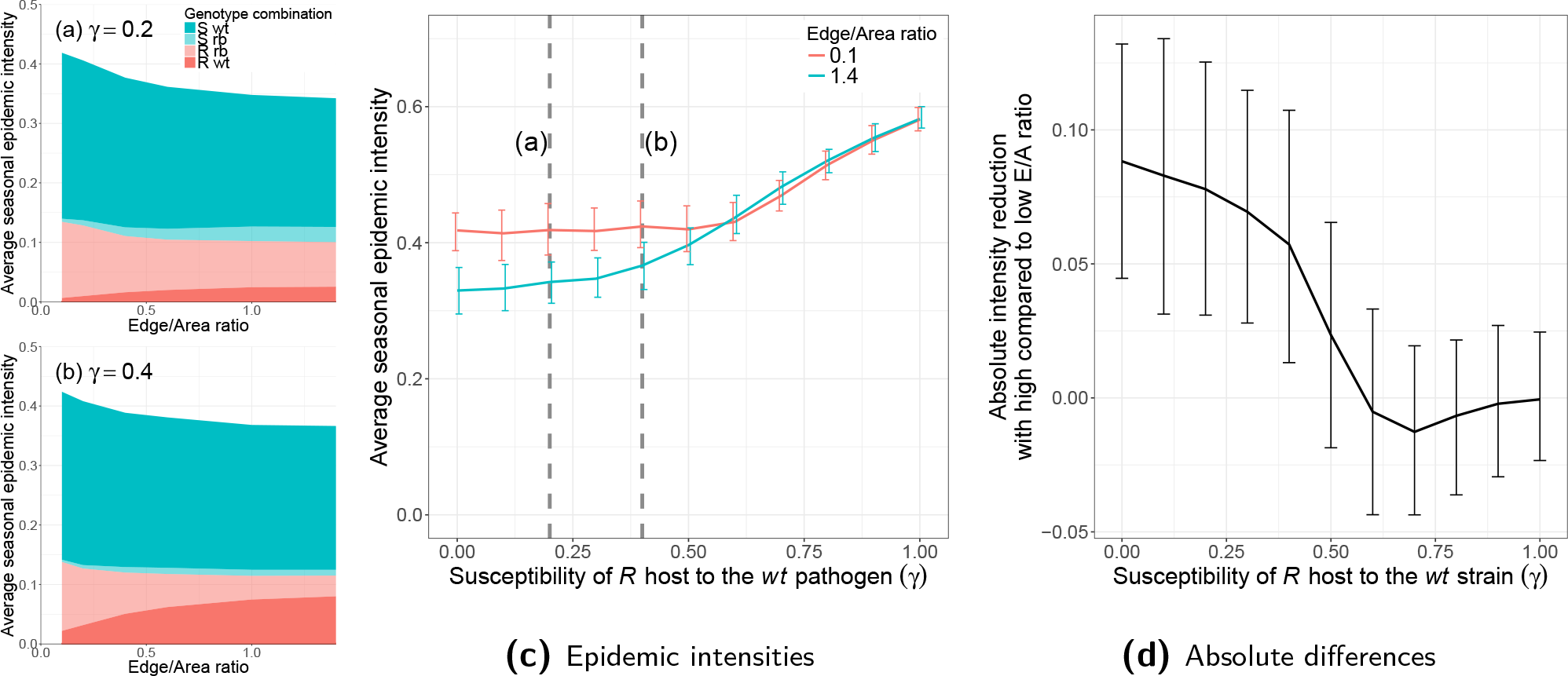
The effect of the susceptibility of the *R* host to the *wt* pathogen strain (*γ*) on the reduction in epidemic intensities from using a high compared to a low landscape edge/area ratio. **The strength of the spatial effect is decreased at lower efficacies of the host *R* gene, as long as the *R* gene is not so weak that the resistance breaking strain is not fit enough to invade the landscape.** The average epidemic intensities resulting from the low and high ends of the E/A ratio scale are shown in (c), while the absolute differences between the results for these two E/A ratio values are shown in (d). Note that using the proportional differences in epidemic intensity produces a qualitatively similar pattern. The relationship between the full range of landscape edge/area ratios and epidemic intensity, for each host/pathogen genotype combination, is shown for two values of *γ* in (a) and (b), and their corresponding positions along the x axis in (c) are marked. Error bars show the 5th and 95th percentiles of the simulation replicates with stochastic landscape generation. For all simulations presented here, the number of seasons *n*_*y*_ = 40, the cost of the *rb* trait *δ* = 0.3 and the kernel parameter *η* = 2.

There is a range of *γ* values, from 0.6 to 0.9 for *δ* = 0.3 (Fig. 6d), for which the strength of the spatial suppressive effect dips below zero and becomes negative, indicating that smaller scales of spatial heterogeneity increase epidemic intensities. This increase only occurs at *δ* and *γ* combinations where the *rb* strain is not fit enough to invade, or is only present at extremely low frequencies (Fig. 7). In addition to this, the increased epidemic intensities are only observed at medium to high values of *γ*, and are most apparent at medium *γ* values. This effect is due to the inability of the *wt* pathogen genotype to sustain epidemics in the *R* host fields without the presence of nearby *S* fields to act as a source of *wt* inoculum. At higher E/A ratios however, where the fields types are mixed at smaller scales of spatial heterogeneity, the closer proximity of *S* field *wt* sources enables greater infection of the *R* field sinks, thereby increasing the overall proportion of infected plants in the landscape, despite the reduction in epidemic intensities on the *S* fields themselves. At very high values of *γ*, the *wt* genotype is fit enough to better sustain epidemics at large scales of spatial heterogeneity (low E/A ratios), so the spatial effect is reduced in strength (and cannot occur at all with *γ* = 1). At low values of *γ*, spatial suppression of epidemic intensities are still observed, despite the non-invasion of the *rb* strain, due to the very low fitness of the *wt* genotype on the *R* host, which severely limits *wt* epidemics in *R* fields at any scale of spatial heterogeneity.

**Figure 7:**
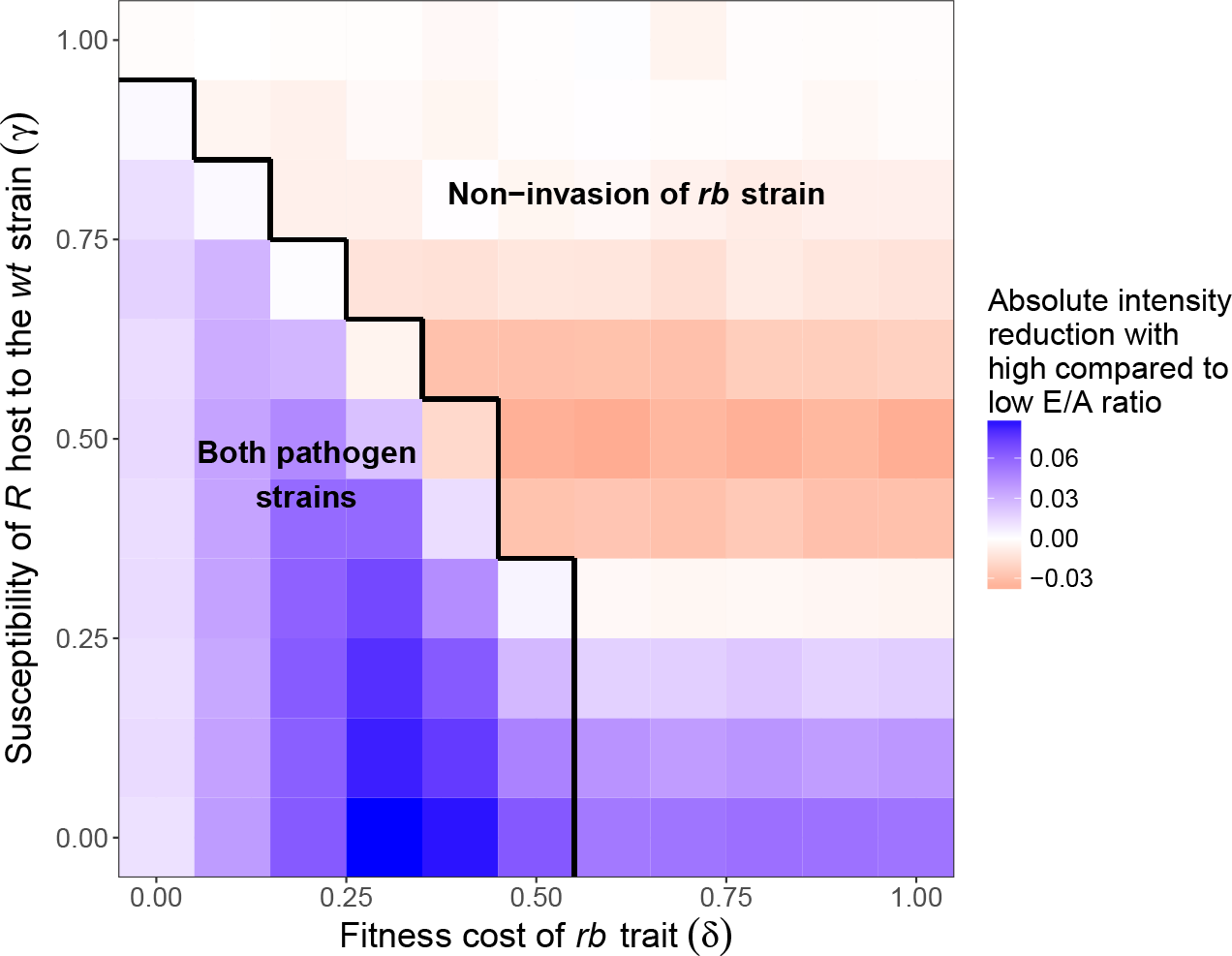
The combined effects of the fitness cost of the resistance breaking trait (*δ*), and the susceptibility of the *R* host to the *wt* pathogen strain (*γ*), on the absolute reduction in epidemic intensities from using a high compared to a low landscape E/A ratio. The baseline epidemic intensities from using a low E/A ratio are shown in Supporting Information Fig. S3. Note that using the proportional differences in epidemic intensity produces a qualitatively similar pattern. **The strongest spatial effect occurs with a 100% effective resistance gene that imposes intermediate fitness costs on the resistance braking pathogen strain. The spatial effect operates in the reverse direction if the resistance breaking strain is not fit enough to invade the landscape, so that only the wild-type strain is present, and the *R* gene is of intermediate to lower efficacy.** In the region where the *rb* strain does not invade (defined arbitrarily as when the *rb* epidemic intensity < 0.01), there is no consistent response to changes in *δ* on the horizontal axis (due to the absence of the *rb* strain). For all simulations presented here, the number of seasons *n*_*y*_ = 40 and the kernel parameter *η* = 2.

### 4.4 Effect of timescale

By relaxing the assumption of a 40 season time period, we can measure the strength of the spatial suppressive effect, or the difference in epidemic intensities between low and high landscape E/A ratios, across a wide range of eco-evolutionary timescales. The average seasonal epidemic intensity generally increases as the number of seasons is increased (Fig. 8a,b,c). This is due to the increasing frequency of the *rb* genotype, which facilitates greater infection of *R* fields as the system approaches its long term evolutionary equilibrium. In the case where the fitness cost of the *rb* trait *δ* = 0, there is no significant spatial suppressive effect on epidemic intensities over 80 seasons (i.e. the red and blue curves converge in Fig. 8a). Here, the *rb* strain is able to completely outcompete the *wt* on both host genotypes at the long term evolutionary equilibrium, meaning that the scale of spatial heterogeneity has no effect. A transient spatial effect does however occur over a low to medium number of seasons, as the system has not yet reached its equilibrium state and the *wt* strain is still present at significant frequencies.

**Figure 8:**
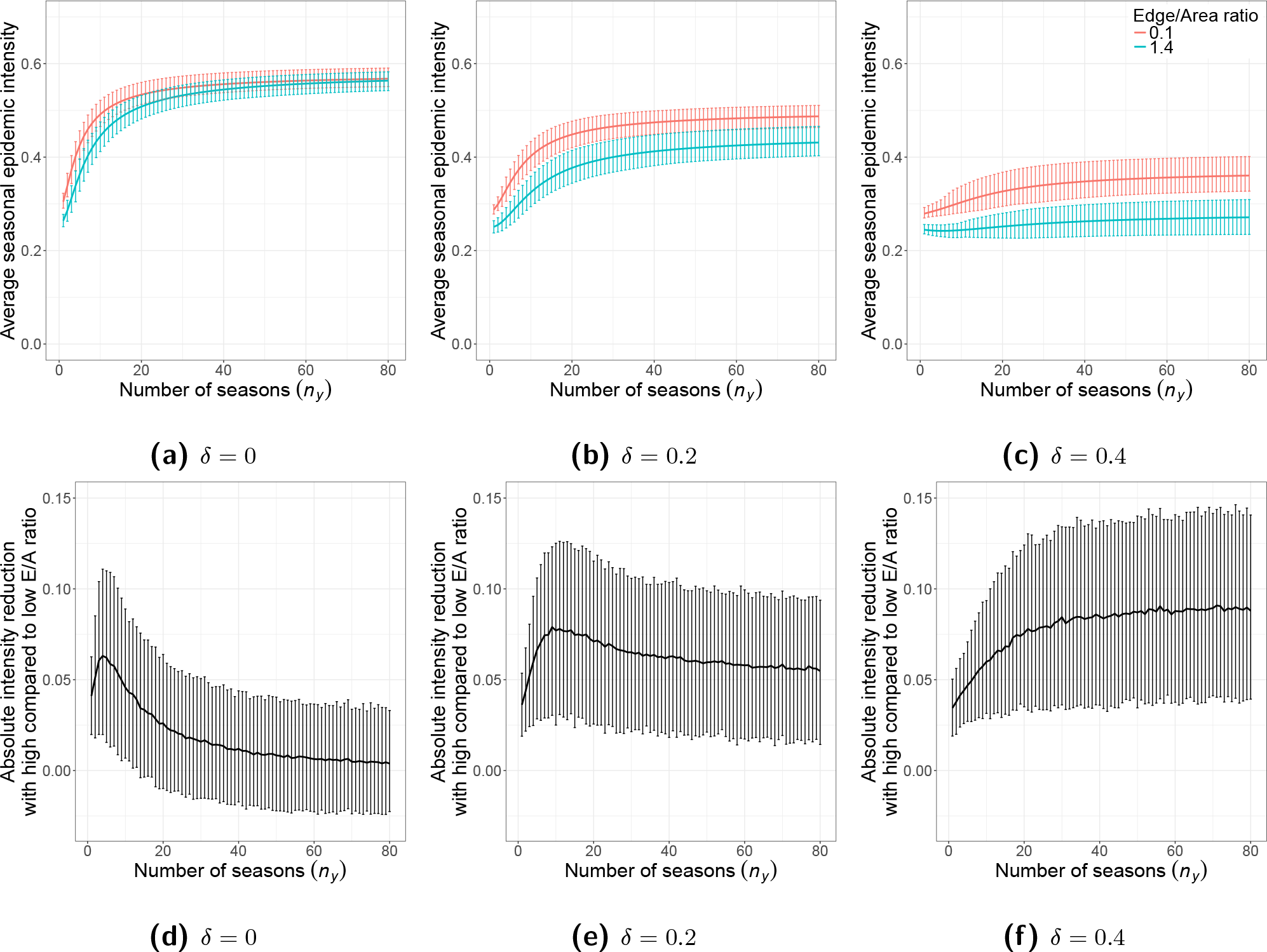
The effect of the number of seasons (*n*_*y*_) on the reduction in epidemic intensities from using a high compared to a low landscape edge/area ratio, at different fitness costs of the resistance breaking trait (*δ*). **The strength of the spatial effect depends on the length of the time frame of interest. There is the potential for a peak effect strength over an intermediate number of seasons, which occurs over a greater number of seasons with a higher fitness cost of the resistance breaking trait.** The average epidemic intensities resulting from the low and high ends of the E/A ratio scale are shown in (a), (b) and (c), while the absolute differences between the results for these two E/A ratio values are shown in (d), (e) and (f). Note that using the proportional differences in epidemic intensity produces a qualitatively similar pattern. Error bars show the 5th and 95th percentiles of the simulation replicates with stochastic landscape generation. For all simulations presented here, the *R* host is completely resistant to the *wt* strain *γ* = 0 and the kernel parameter *η* = 2.

There is a specific number of seasons, for a given fitness cost of the *rb* trait (*δ*), over which the greatest spatial suppressive effect on epidemic intensities can be achieved (Fig. 8d,e,f). For *δ* = 0 and *δ* = 0.2 (Fig. 8d,e) this peak effect strength occurs over a relatively short time period, approximately 4 and 11 seasons respectively, whereas with *δ* = 0.4 the greatest spatial suppressive effect is observed as the system approaches its long term evolutionary equilibrium.

The initial increase in the strength of the spatial suppressive effect, from the first season to the intermediate peak, is due to a faster increase in epidemic intensities at a low E/A ratio compared with a high E/A ratio. This is because the newly emergent *rb* genotype is able to propagate rapidly in the highly aggregated *R* fields. The *rb* genotype initially spreads less quickly in *S* fields, and therefore at high E/A ratios, due to the high level of competition with the coexisting *wt* pathogen genotype. Beyond the peak in spatial suppression, if it is present, the strength of the suppressive effect declines as epidemic intensities begin to increase faster at high compared to low E/A ratios. This is due to the increased importance of the *rb* strain spread on *S* hosts to the sensitivity of the system to changes in season number. The overall *rb* frequency in the landscape is higher, than it is over a lower number of seasons, resulting in a faster growth rate on the *S* host as it competes more effectively with the *wt* strain. The increase in the intensity of the *S rb* epidemic increases the overall connectivity for the *rb* strain in landscapes with a high E/A ratio, thereby increasing the epidemic intensity for the *R rb* epidemic as well. The *rb* spread on the *R* host with a low E/A ratio however, is already well advanced at this stage, causing its rate of spread to decline.

The pattern here is that of a trade-off in the relative sensitivity to change in season number of the *rb* epidemics in the *S* and *R* hosts, with the peak in the strength of spatial suppression being where the combined sensitivity effect of the two processes is the same at both small and large scales of spatial heterogeneity. This peak occurs over a higher number of seasons with larger values of *δ* because the greater trait cost means that the *rb* strain spreads more slowly on the *R* host, and takes longer to become competitive with the *wt* strain on the *S* host. There is no intermediate peak with *δ* = 0.4 (Fig. 8f) because the *rb* trait is too costly for the *rb* strain to spread as effectively on the *S* host (Fig. 4c).

The variation in epidemic intensities due to the stochastic placement of fields in the landscape (size of the error bars), is highest at the points where the strength of the spatial suppressive effect is greatest (Fig. 8). This implies that the effect of the specific stochastic arrangement of fields, and therefore the precise degree of spatial heterogeneity, is greatest when the general sensitivity to spatial dynamics is maximised.

## 5 Discussion

Our model shows that planting susceptible and resistant host crop fields at smaller scales of spatial heterogeneity reduces epidemic intensities over a wide range of eco-evolutionary timescales. Such spatial strategies would therefore increase the durability of disease resistance, using a definition of durability analogous to the additional yield measured by the number of uninfected host growth days (van den Bosch and Gilligan, 2003). The underlying mechanism is similar to the dilution effect that has been reported to reduce short term epidemic intensities in within field cultivar mixtures (Mundt, 2002). Smaller scales of spatial heterogeneity reduce the local density of a given crop cultivar, meaning that some of a pathogen strain’s potential force of infection is wasted as its inoculum disperses onto other nearby cultivars on which it has a lower reproductive fitness. This ‘wasted inoculum’ effect generally suppresses the epidemics caused by each pathogen strain on the host cultivar where they are specialised and have the greatest fitness, while to a smaller extent boosting epidemics of that strain on their less preferred host. Whilst this study has focussed on qualitative plant resistance genes in a gene-for-gene system, these dynamics should be generally applicable to any system where multiple pathogen strains have different relative fitnesses when infecting multiple different host genotypes. Indeed, a similar basic spatial effect on resistance durability was observed by Papaïx et al. (2018), in a model in which the resistance trait gradually eroded due to progressive small mutations in the pathogen population.

Our results also shed light on a number of the factors determining the strength of this spatial suppressive effect. This has significant implications for the potential effectiveness of any such spatial strategy if it were implemented by growers, as the benefit in terms of crop yield must outweigh any potential economic costs of farming in such a manner. Growing monoculture crops in very large fields, with consequentially large scales of spatial heterogeneity, is the norm in many systems of developed agriculture, largely for reasons of economic efficiency. It is therefore important that we characterise the fundamental eco-evolutionary processes that interact with host spatial structure, as these will ultimately play a crucial role in identifying the specific pathosystems where such spatial strategies are most likely to succeed.

The scale of pathogen dispersal within the agricultural landscape must correspond in some sense to the scale of heterogeneity implemented in that landscape in order to maximise the effectiveness of a spatial diversification strategy. Given that lower mean dispersal distances lead to the spatial suppressive effect being relevant over a wider range of scales (Fig. 3), pathogens with more restricted ranges of landscape scale dispersal are more likely to be effectively controlled in this manner. The underlying idea of requiring different spatial diversification strategies to control pathogens with different dispersal characteristics is supported by Sapoukhina et al. (2010). That work, however, focussed on the qualitative differences between local short ranged dispersal through diffusion, and stratified dispersal that also included a separate long distance component, rather than looking at spatial heterogeneity as a continuous scale.

The spatial suppressive effect of cropping pattern on epidemic spread is maximised at an intermediate value for the fitness cost associated with the resistance breaking trait (Fig. 5). This peak in effect strength is ultimately driven by landscapes with different scales of spatial heterogeneity generating different frequencies of interaction between the various host and pathogen genotypes in the system. The epidemic intensities for these different infection classes respond at different rates to changes in the cost of the resistance breaking trait, depending on the current value of that trait (Supporting Information Fig. S2). It is these different rates of change that drive the variable strength of the spatial suppressive effect, and create the peak effect strength at intermediate fitness cost values. The exact fitness cost value for this peak in the spatial suppressive effect is lower with a less effective resistance gene (Fig. 7).

A less effective resistance gene lowers the strength of the spatial effect, as long as the resistance breaking strain has a high enough fitness to be able to invade the agricultural landscape (Figs 6, 7). If this is not the case, and only the wild-type strain is present, intermediate to lower efficacy *R* genes can drive a reverse spatial suppressive effect, where epidemic intensities are higher with smaller scales of spatial heterogeneity. This occurs because wild-type epidemics are only able to sustain themselves in partially resistant fields when there are nearby susceptible fields to act as inoculum sources. The effect is similar to that observed by Papaïx et al. (2014), who showed in a single strain system that the directional effect of spatial aggregation depended on the *R*_0_ value for the disease epidemic on the resistant variety.

The combined genetic context in this system is created by the combination of the fitness cost of the resistance breaking trait and the efficacy of the resistance gene. From this we can conclude that a spatial diversification strategy is most likely to be cost effective when using a 100% effective major resistance gene that imposes intermediate fitness costs on a resistance breaking pathogen strain. Despite this, a spatial strategy is still likely to be at least partially effective in any genetic context, as long as the resistance breaking pathogen strain is fit enough to invade and persist within the landscape (Fig. 7). The fact that spatial diversification can actually worsen epidemics when only a wild-type strain is present, and a partially effective resistance gene is used, highlights the necessity of understanding the state of the pathogen community and the genetic nature of the system before implementing such control strategies. A spatial strategy will be less effective when there are no fitness costs for the resistance breaking strain, however an effect is still observed due to the time required for this strain to fully take over the pathogen population. This naturally becomes particularly apparent when looking over a lower number of seasons.

A critical factor that is generally neglected within the study of resistance durability is the length of the time frame of interest. If we consider any improvement in durability to be the yield gain achieved, this will naturally depend on the time period over which we measure such gains, which in practice should itself depend on factors such as the frequency with which new resistant cultivars are developed. This timescale will obviously vary for different crop disease systems, and will play a significant role in determining whether a spatial strategy has a large enough effect to be economical for practical use. The strength of the spatial suppressive effect depends on the number of seasons over which it is measured, with the potential for a peak in spatial effect strength over an intermediate number of seasons (Fig. 8), i.e. in between a short term epidemiological timescale and the long term evolutionary equilibrium. In a similar manner to the effect of the fitness cost of the resistance breaking trait, this occurs because the different infection classes in the system respond at different and varying rates to changes in season number. The frequency of these host-pathogen genotype combinations, and therefore the effect they have on overall epidemic intensity, depends on the scale of spatial heterogeneity in the distribution of host genotypes within the landscape. The resultant peak spatial effect occurs over a higher number of seasons with a higher cost of the resistance breaking trait, due to the slower spread rate of this resistance breaking strain. Generally this means that a spatial strategy is most likely to be effective over short timescales for resistance breaking strains that carry little or no fitness costs, and over longer timescales for more costly traits.

In the current study we have restricted the cropping ratio of the susceptible and resistant cultivars to 50 : 50, in order to avoid having to consider potential interactions between the effects of the scale of spatial heterogeneity and the amount of resistant crop deployed. The potential ways in which the patterns we have described might be influenced by different cropping ratios is a valid area for further study however, as is the way that optimal cropping ratios might in turn be influenced by spatial dynamics. The non-spatial model of Fabre et al. (2012) demonstrated that the optimal cropping ratio (i.e. the proportion of resistant fields) varied from intermediate to high values, and depended among other factors on the relative contributions of within field, between field and reservoir driven infection. Demographic stochasticity, which has been shown to bias optimal cropping ratios towards higher values, is another potential route for further investigation (Lo Iacono et al., 2013). The associated chance of pathogen strain extinction at low frequencies or under periodic perturbation could potentially interact with the effects of patch size and spatial heterogeneity on disease dynamics.

In conclusion, this study has demonstrated the key effect that spatial structure can have on disease resistance durability. The diversification of resistance genes at small scales of spatial heterogeneity is a potentially valuable strategy for improving long term crop yields, depending on whether the strength of the spatial effect leads to such a strategy being economical. Factors such as the pathogen dispersal scale, the genetic properties of the host-pathogen interaction, and the time frame of interest play a crucial role, and highlight the need for a thorough understanding of any disease system to which this strategy is applied.

## 6 Acknowledgements

We thank Elliott Bussell, Richard Stutt, Andrew Craig and Eleftherios Avramidis for helpful discussions and technical advice related to this project. B. Watkinson-Powell acknowledges the Biotechnology and Biological Sciences Research Council of the United Kingdom (BBSRC) for support, via a University of Cambridge DTP PhD studentship.

