## Supporting Information: Notes and Figures for "When does spatial diversification usefully maximise the durability of crop disease resistance?"

#### Notes S1: Fully susceptible model

A landscape made up of only susceptible fields forms the baseline epidemiological context to which all other simulations (with resistance included) are compared. In this model the only state variable of interest is $I_{S,x,y}$, the number of infected plants in a representative $S$ field in season $y$. Healthy plants can become infected through three alternative routes: from the $I_{S,x,y}$ infected plants in the same field at rate $\beta_F$, from infected plants in other fields at rate $\beta_C$, and from the reservoir at rate $\alpha_E$. When combined these rates give the ODE for the change in the number of infected plants as:

$$\frac{\mathrm{d}I_{S,x,y}}{\mathrm{d}t} = (n_p - I_{S,x,y})(\alpha_E + \beta_C \left( \sum_{z=1}^{n_f} I_{S,z,y} K[z,x] \right) + \beta_F I_{S,x,y}) \tag{1}$$

where $x$ indicates variables pertaining to a particular field, $K[z,x]$ is the dispersal kernel coupling field $z$ to field $x$, and $z \neq x$. The dispersal kernel follows a normalised negative exponential distribution of the form $K = \frac{\eta^2}{2\pi} e^{-\eta d}$. In all simulations, $I_{S,x,y}(0) = 0$ (i.e. the number of infected plants within fields is set to zero at the beginning of each season), with all epidemics started by infection from primary reservoir inoculum. Solving Eqn. (1) numerically, then integrating with respect to time, gives the area under the disease progress curve (AUDPC). The AUDPC for this baseline model (where the proportion of $R$ fields $\phi = 0$) is $A_0 = \sum_{x=1}^{n_f} (\int_0^{n_d} I_{S,x,y}(t)\,\mathrm{d}t)$, where $n_f$ is the number of fields and $n_d$ is the number of days in a season. This is used for comparison with later simulations to measure the reduction in epidemic intensity with the inclusion of resistant fields and the resistance breaking pathogen. The epidemic behaviour and intensity in this fully susceptible model does not change between seasons as there is no transformation in the composition of the reservoir component over time. In the full model from the main paper on the other hand, the rates of primary infection from the reservoirs, derived from the constant $\alpha_E$ value and given by $\alpha_{wt,x,y}$ and $\alpha_{rb,x,y}$, do change over time.

The values of the parameters $\beta_F$, $\beta_C$ and $\alpha_E$ used in the simulations were determined by calculating the relative contributions of the three infection routes to, and the intensity of, the overall landscape epidemic. The overall intensity of the epidemic, which describes the proportion of infected plants, averaged over a season, is given by $\Omega_{int} = A_0/(n_f n_p n_d)$. The relative contribution of each route of infection is described by the 'epidemic profile', which is given by $\Omega_{pfl} = (\Omega^1_{pfl}, 1 - \Omega^1_{pfl} - \Omega^3_{pfl}, \Omega^3_{pfl})$, where the three components are the relative contributions of the reservoir, the between field contacts, and the within field contacts respectively.

The following equations are used to calculate the contributions of the reservoir (Eqn. (2)) and within field infection (Eqn. (3)) at time $t$.

$$\frac{\mathrm{d}I^1_{\bar{S},y}}{\mathrm{d}t} = (n_p - I_{\bar{S},y}(t))\alpha_E \tag{2}$$

$$\frac{\mathrm{d}I^3_{\bar{S},y}}{\mathrm{d}t} = (n_p - I_{\bar{S},y}(t))\beta_F I_{\bar{S},y}(t) \tag{3}$$

Here, $I_{\bar{S},y}(t)$ is the average number of infected plants per field in the spatially explicit baseline model (Eqn. (1)) at time $t$. Taking the AUDPCs of Eqns (2) and (3), we calculate $\Omega^1_{pfl} = \frac{n_f}{A_0}\int_0^{n_d} I^1_{\bar{S},y}(t)\,\mathrm{d}t$ and $\Omega^3_{pfl} = \frac{n_f}{A_0}\int_0^{n_d} I^3_{\bar{S},y}(t)\,\mathrm{d}t$, which gives the proportion of the total number of infected plants that are infected by the reservoir and by within field transmission respectively. The proportion of infected plants that are infected by between field transmission is calculated by taking the remaining proportion of infected plants that did not become infected by one of the other two routes of transmission ($\Omega^2_{pfl} = 1 - \Omega^1_{pfl} - \Omega^3_{pfl}$). This is done because the value of $\Omega^2_{pfl}$ is more difficult to compute directly as it involves spatially explicit interactions that cannot be derived from average field outcomes as in Eqns (2) and (3).

The R optimiser function *nlminb* was used to find the $\beta_F$, $\beta_C$ and $\alpha_E$ values that gave $\Omega_{int} = 0.5$ and $\Omega_{pfl} = (1/3, 1/3, 1/3)$. The objective function for this optimisation is given by
$E = (\Omega_{int} - 0.5)^2 + (\Omega^1_{pfl} - 0.333)^2 + (\Omega^2_{pfl} - 0.333)^2$, where $\Omega^2_{pfl} = 1 - \Omega^1_{pfl} - \Omega^3_{pfl}$. In this way, for a fully susceptible model, the three transmission routes are of equal importance in maintaining the epidemic, and half of the plants in the landscape become infected over a season.

Optimising the model to set infection rate parameters for every random replicate set of field coordinates is computationally expensive, and undesirable for the purposes of reproducibility. Therefore, the fields in the fully susceptible model used to optimise the parameters and provide a baseline epidemic scenario were arranged in a regular square grid pattern (with 1 arbitrary distance unit gaps between fields) within the $10$x$10$ landscape. This of course means that the random landscape used for each simulation replicate produces a slightly different epidemic intensity and profile. This is of limited concern however, as the purpose of the epidemic parameter optimisation is merely to create epidemics of a general size and balanced structure that allow convenient manipulation and comparison.

#### Notes S2: Interior landscape Edge/Area ratio

The interior edge/area ratio of a landscape is calculated by summing the total length of the edges separating different patch types, and dividing by the total area of the landscape. For example in a $10x10$ landscape divided into two halves, by a line perpendicular to the landscape perimeter (Fig. S1), the interior edge is of length 10, and so the edge/area ratio is $10/10^2 = 0.1$.

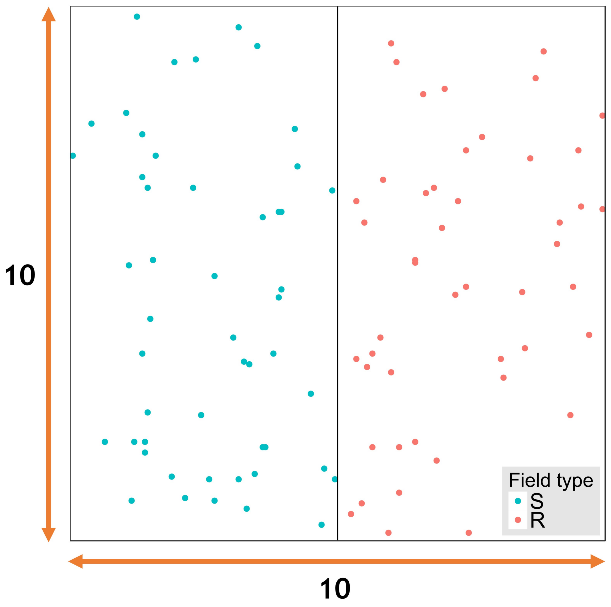


**Figure S1:** Example random field arrangement for a template that divides the agricultural landscape into two patches. The labelled dimensions of the landscape template are used to calculate the edge/area ratio $(10/10^2 = 0.1)$, where the numerator of the fraction is the length of the internal edge $(= 10)$ and the denominator is the area $(= 10^2)$.

The patches created by the landscape template are categorised as being either of type $S$ or type $R$ in an alternating pattern. In each replicate simulation, individual fields are then designated (according to the cropping ratio $\phi$) as being either $S$ or $R$, and are then assigned a location with random coordinates drawn from a uniform distribution within the patches pertaining to that genotype.

### Additional figures

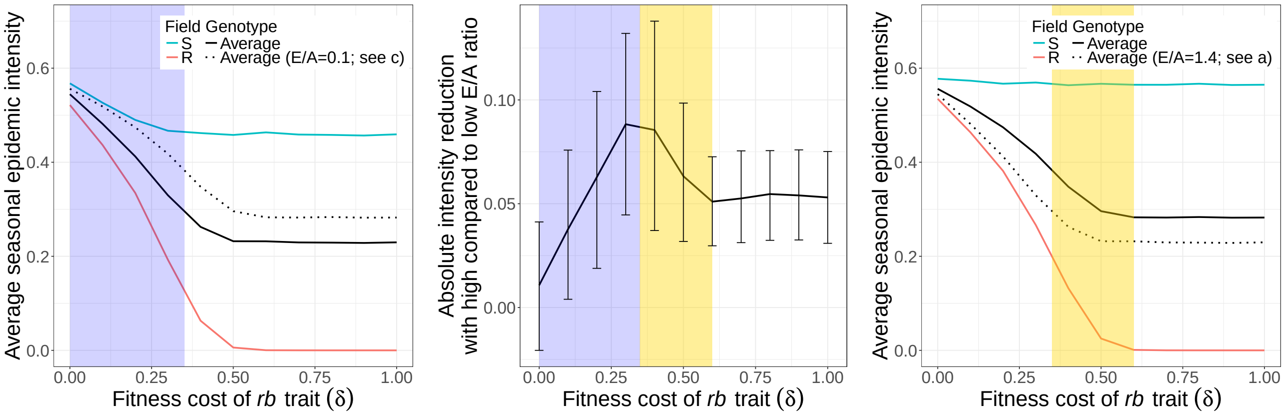


**(a)** Edge/Area ratio $= 1.4$ **(b)** Absolute differences **(c)** Edge/Area ratio $= 0.1$

**Figure S2:** The effect of the fitness cost of the resistance breaking trait ($\delta$) on the reduction in epidemic intensities from using a high compared to a low landscape edge/area ratio. The average epidemic intensities, separated into field types (coloured lines), resulting from a high E/A ratio are shown in (a), and from a low E/A ratio in (c). The total epidemic intensities across both field types (i.e. the mean of the red and blue responses; also shown in main text Fig. 5a) are given as a solid black line for the E/A ratios in (a) and (c). The total epidemic intensities for the opposite ends of the E/A ratio scale are also included as dotted lines in (a) and (c). The absolute differences between the total epidemic intensities for these two E/A ratio values are shown in (b). Note that using the proportional differences in epidemic intensity produces a qualitatively similar pattern. Error bars show the 5th and 95th percentiles of the simulation replicates with stochastic landscape generation. **In the blue shaded region, the system changes more rapidly with $\delta$ at a high E/A ratio (a), due to the activity of the $rb$ pathogen on the $S$ host (blue line in (a)). Beyond the intermediate peak in (b), in the gold shaded region, the system changes more rapidly with $\delta$ at a low E/A ratio (c), due to the activity of the $rb$ pathogen on the $R$ host (red line in (c)).** For all simulations presented here, the number of seasons $n_y = 40$, the $R$ host is completely resistant to the $wt$ strain $\gamma = 0$ and the kernel parameter $\eta = 2$.

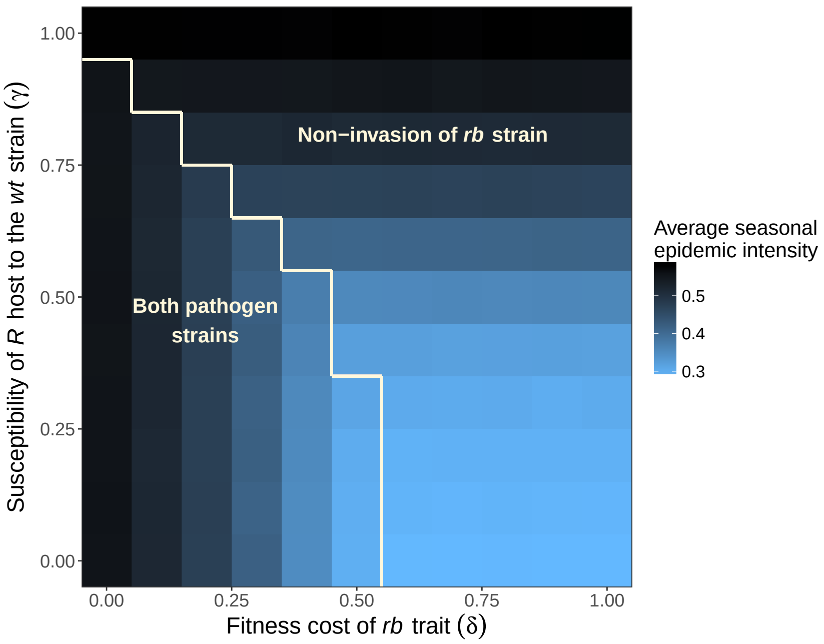


**Figure S3:** The combined effects of the fitness cost of the resistance breaking trait ($\delta$), and the susceptibility of the $R$ host to the $wt$ pathogen strain ($\gamma$), on the average epidemic intensities resulting from a low E/A ratio $= 0.1$. This plot serves as a baseline for the absolute reduction in epidemic intensities from using a high compared to a low E/A ratio (main text Fig. 7). In the region where the $rb$ strain does not invade (defined arbitrarily as when the $rb$ epidemic intensity $< 0.01$), there is no consistent response to changes in $\delta$ on the horizontal axis (due to the absence of the $rb$ strain). For all simulations presented here, the number of seasons $n_y = 40$ and the kernel parameter $\eta = 2$.
